# The Testing and Characterization of the Bacterial Response to Novel Lipid Ether Amine-Based Antibiotics

**DOI:** 10.64898/2026.07.22.740093

**Authors:** Daniel J. Gibson, Artem S. Svetlov

## Abstract

Bacterial biofilms exhibit enhanced resistance to antibiotics compared to planktonic bacteria, though the mechanisms underlying this tolerance remain incompletely understood. Evidence suggests biofilm-associated bacteria maintain active lipid metabolism despite reduced metabolic rates, making lipid-targeting strategies potentially effective. We investigated isopropyl lipid ether amines (LEAs), computationally designed to bind phospholipase A2, as novel antibiotics targeting biofilm bacteria through lipid metabolism disruption. LEAs consist of variable-length alkyl chain analogs of natural fatty acids connected via ether linkages to cationic head groups. Using methicillin-resistant Staphylococcus aureus (MRSA) and Pseudomonas aeruginosa in collagen microplate assays, minimal inhibitory concentration studies, and ex vivo porcine biofilm models, we systematically compared LEAs against their fatty acid analogs to isolate the contribution of the lipid and amine moieties. Palmitic acid analog LEA-160 and oleic acid analog LEA-181 achieved MIC and reduced microplate biofilm colony forming units (CFU) against MRSA, while short lipid chain octanoic acid analog LEA-80 reduced Pseudomonas biofilms, with no effect for natural fatty acids. This demonstrates that the cationic amine group provides essential antibacterial function beyond the alkyl chain contribution. Comprehensive lipidomics analysis using LC-IMS-MS/MS revealed that LEA treatment induces significant alterations in MRSA lipid profiles, supporting a mechanism involving disruption of bacterial membrane lipid metabolism. Structure-activity relationships confirmed that both the lipid chain and cationic moieties are necessary for LEA antibacterial efficacy, with metabolic effects distinct from their natural fatty acid analogs. These findings establish LEAs as a mechanistically distinct antibiotic class targeting bacterial lipid metabolism pathways critical for biofilm survival.

## Introduction

Surface aggregation of bacteria into biofilms had been underappreciated in many fields of medicine^[1]^. However, their resiliency in the face of modern antibiotics^[2]^ and their role in implanted device complications^[3-7]^ have led to a new understanding of their role in chronic diseases. There are still very few techniques for studying medically relevant biofilms and consequently very few treatments have emerged; most of which require physical removal of the affected tissue or device. Systemic treatments, like antibiotics, are direly needed.

Current efforts to generate biofilm-targeted therapies aim at the biofilm’s connection to the surface, inhibition of proliferation and spread, or eradication.^[8]^ For surfaces that are not readily accessible, dispersal agents informed either by physical chemistry or quorum sensing mechanisms are the most popular. However, in medical contexts, solely dispersing the biofilm would be dangerous, as it would allow for systemic spread of the bacteria. New antibiotics which can inhibit or kill the biofilm-associated bacteria are an anticipated necessity prior to the physical removal or resorption of the biofilm mass.

Evidence from a few studies suggest that while biofilm-associated bacteria are less metabolically active than planktonic, they are not completely dormant. Even if there is a high degree of dormancy, the molecular complexes responsible for exiting dormancy may serve as potential targets. Empirically, biofilms have been found to be more lipid-rich than equivalent counts of planktonic bacterial cells,^[9]^ indicating that lipid metabolism may be one targetable process. However, daptomycin and polymyxins are current antibiotics which target lipid membranes and some resistance has been discovered.^[10, 11]^ Targets that are more upstream from the membrane itself may be necessary.

Evidence of current antibiotics activities against biofilms exists but is incomplete. It is often stated that current antibiotics do not kill biofilm-associated bacteria, however, some evidence from lab studies suggest that most of the bacterial cells within the biofilm mass are still susceptible. In the ex vivo porcine skin wound model, the bacteria are grown on the denuded skin surface in the absence of a liquid solution. Based on scanning electron microscopy, attached bacteria are the predominantly observed cells,^[12]^ but 3-4 log_10_ of these bacteria do die in the presence of 50× MIC antibiotic^[13]^. This susceptibility of most of the bacteria in a biofilm has long been known.^[1]^ Studies with antiseptics evidenced that the agents can still kill the bacteria in a biofilm, but that they may need a longer treatment time.^[2]^ The remaining bacteria have been called either “persister cells” or “functional biofilm”, with the latter being derived from the sensitive bacteria that remain after antibiotic treatment. While current antibiotics can kill some biofilm-associated bacteria, they are not effective enough. The use of biofilm-based models earlier in the design and testing phase is expected to be necessary to generate biofilm-eradicating antibiotics.

Herein we describe the initial testing of isopropyl lipid ether amines (LEAs) in the ability to kill sessile biofilm-associated bacteria. These molecules were initially designed in silico for their ability to bind phospholipase A2. Each LEA has an alkyl chain of varying length linked to a cationic head group through an ether bond. The LEAs were tested side-by-side with the alkyl-equivalent fatty acid in an attempt to demonstrate the contributions of the alkyl chain. In addition to cell viability assays, initial skin toleration studies and a bacterial secreted lipidomics study were completed. We present evidence that both the alkyl chain and the amine-containing moiety are required for effectiveness and that they lead to a differential lipid metabolism response in methicillin resistant *Staphylococcus aureus*.

## Experimental Section

### Bacterial Stocks & Inoculum Preparation

Bacterial stocks were sourced from ATCC. For Gram positive, a methicillin-resistant strain of *Staphylococcus aureus* (SA, BAA44) and methicillin-sensitive strain SA 29213 were used, while for Gram negative, *Pseudomonas aeruginosa* (PAO1) was used.

New cultures were started by sampling a small portion of the partially thawed glycerol stock and inoculating 2.0mL of tryptic soy broth in a sterile 15mL conical tube. The tube was sealed and agitated in an incubator set to 35°C overnight. The overnight broth was sampled and bacterial count estimation was done by turbidimetric methods.

### Collagen Microplate Assay

Early log-phase tryptic soy broth (TSB) culture stocks of each species/strain were seeded in collagen I coated 96-well plates at 5×10^5^ CFU/well (T_0_, n = 8). For adherent SA 29213 and MRSA BAA-44 cultures, after 24h (T_1_) the culture broth was discarded, the wells were gently rinsed with PBS-Tween (PBS-T) to remove residual planktonic bacteria, and replaced with fresh TSB. Biofilm rafts formed at the air/fluid interface in PAO1 cultures were removed from each well using microforceps, gently rinsed in a plate with PBS-T, then transferred to a separate plate with TSB. After further 48h of incubation (T_2_), pre-treatment baseline samples were collected and the treatment media (T_3_) was added. Following another 24h of treatment (T_4_), wells with adherent cultures were washed with PBS-T and both adherent and PAO1 biofilm cultures vigorously pipetted to disaggregate biofilms. The samples were plated from 10^-3^ to 10^-5^ using an automatic spiral plater (Interscience easySpiral Dilute, Interscience, Saint-Nom-la-Bretèche, France), and automatically counted for CFU (Interscience Scan 300, Interscience, Saint-Nom-la-Bretèche, France).

Log-logistic dose-response curves were fitted using R drc library, with log(ED50) as a fitting parameter. Where a lower limit parameter could be fit, 4- or 5-parameter regression was used, otherwise a 3-parameter curve was fit with the lower limit set to 0. Effective doses and 95% confidence intervals were calculated using the ED.drc function.

### Computational ligand-docking screen

To identify antimicrobial mechanisms for LEA compounds, an end-to-end protein-ligand docking pipeline was developed to procedurally generate flexible conformers for LEA analogs with varied lipid chain length (8-20 carbon atoms) and saturation (at 0-4 sites, cis and trans), endogenous phospholipid and fatty acid ligands, and a propranolol decoy control matching the LEA polar head moiety. Using crystallographic structures of 34 putative targets, predominantly lipid metabolic enzymes, virulence factors, receptors, and transcriptional regulators, docking poses were analyzed and binding affinities were predicted using AutoDock VINA. Binding poses were optimized for change in free energy of the bound vs unbound conformations (ΔG, kcal/mol), each -1.0 change in ΔG indicating 10× higher affinity. Catalytic binding sites were further refined for 10 secreted bacterial virulence factors and 7 endogenous human phospholipases (Table S1). Binding affinity was then exhaustively sampled with 8 replicates for each LEA or endogenous ligand and affinity data across all targets was analyzed using a gaussian Generalized Linear Model (GLM) with the statsmodels python library. A global affinity model, with individual receptor and pooled endogenous ligand terms (0 for LEA compounds, 1 for endogenous ligands), was used to evaluate binding affinity of LEA compounds vs endogenous ligands over-all. Ligand-specific effects were additionally modeled using individual ligand, receptor, pooled endogenous ligand, and receptor-by-endogenous ligand interaction terms. Relative binding affinity of LEA vs each target’s endogenous ligand(s) was used to identify optimized LEA candidates.

### In-vitro Skin Construct Tolerability

A full thickness in vitro human skin model (MatTek EpiDermFT) was treated with topically formulated LEA compounds to evaluate skin tolerability markers LDH and IL-1β. The 1cm skin constructs were provided pre-wounded using a 3mm biopsy punch, cultured, and treated in 5mL serum-free media according to manufacturer specifications. Topical nanoparticle formulations of 500μM and 1000μM LEA compounds in 20% Pluronic-F12 PBS were prepared using thin film hydration, ultrasonication, and unilamellar vesicle extrusion with 100nm pore size (Mini-Extruder, Avanti Lipids). Following equilibration of 6-well culture inserts for 24hr in EpiDermFT Assay Medium, media was sampled for assays, exchanged for fresh Assay Medium, and treated with formulated LEA compounds or vehicle control by applying 100μl of the test compounds to the surface of the skin constructs. Media was sampled daily, exchanged after 72hr, and treated with an additional 100μL of test compounds for 48hr. Following 6 days in culture, histopathology slides were prepared and stained using H&E to evaluate viability and epithelial closure of the simulated wound.

Media samples were analyzed using IL-1β ELISA (Invitrogen) and LDH (Thermo-Fisher CyQUANT-LDH) assays following manufacturer specifications and timepoints were evaluated using repeated measures ANOVA.

### Minimal Inhibitory Concentration Studies

In order to determine the initial dosing and activity versus the controls, a 96-well minimal inhibitory concentration assay was run per ISO 20776-1:2019. Stocks for each item were created at 250mM with DMSO and subsequent dilutions were made to maintain the DMSO at 1% for each test well.

### Ex-Vivo Porcine Skin Wound Model

Food-grade pre-processed porcine skin was obtained from an on-line food supplier. The skins were separated and placed individually in freezer-grade food storage bags and allowed to freeze in a flattened layout. A sheet of frozen skin was removed from the freezer and circular coupons were cut from the sheet using a 12 mm punch-press. A rotary tool ball-shaped burr was used to create a wound in the center of each coupon. The coupons were then placed in to a 50mL tube (10 per tube), and were rinsed and sterilized by an alternating cycle of 10% bleach and 70% ethanol as previously reported^[13]^. After a final series of rinsing, the coupons were stored in sterile phosphate buffered saline (PBS) and were used within 24 hr.

Non-supplemented soft agar was prepared to 1.2% (w/v) final concentration in sterile water and autoclaved to melt and sterilize the agar. The agar was allowed to cool to ∼40°C prior to pouring into 90 mm petri dishes. The dishes were allowed to solidify and cool over-night, re-stacked and wrapped, and then stored at 4°C until needed.

Sterilized explants were placed on soft agar (6 per plate) and both were allowed to come to room temperature. The wounds were inoculated with 10 μl of diluted overnight cultures at 10^6^ CFU per coupon. The explants were covered with sterile saline moistened gauze and a sterile microscope slide was placed on top to hold the gauze in place. The plates were then placed into an incubator at 35°C for 3 days for mature biofilm formation. Each explant was then removed and placed into a single well in a sterile 12-well plate. Antibiotics were added to each well, oxacillin for BAA44 and gentamicin for PAO1 at greater than 50× MIC as previously reported^[13]^ and allowed to incubate overnight at 35°C. The explants were rinsed 3× with PBS-T, and the first batch of explants were harvested as time point 0 for bacterial enumeration. The remaining explants were subjected to exposure to the test items or controls overnight at 35°C. The remaining explants were harvested for bacterial CFU quantification.

### Test Agents & Controls

For testing, three LEAs were utilized. The sole difference among the LEA were the alkyl chain length and degree and state of saturation. LEA compounds are synthesized from their respective fatty alcohols by formation of an epoxide intermediate with epichlorohydrin, followed by nucleophilic substitution by isopropylamine (Alchem Laboratories Corporation). For controls, 3 organic fatty acids with the same lipid chain were used in order to control for effects solely attributable to the lipid moiety, i.e. octanoic fatty acid for LEA-80, oleic acid for LEA-181, and arachidonic acid for LEA-204. Additionally, vancomycin (a currently popular antibiotic) and 10% bleach (the standard for decontamination) were used to assess how current modalities and standards behave under the same test conditions.

### Lipidomics Data Collection

Lipids were extracted from the treated bacterial cultures and media using two phase separation and drying of the organic phase using 2:1 MeOH/Chloroform. Samples were then reconstituted in the glass that they were received in. Initially 200 μl 90/10 (v/v) methanol/chloroform was added to the tubes. Samples were sonicated in two batches for 10 min/batch. Solvent was transferred to 1.5mL glass total recovery vials. Tubes appearing to have residual residue had an additional 100μL chloroform (100%) added. After washing, this volume was then combined with the previous material for a total volume of 300 uL. A portion of these samples were then analyzed to determine total lipid concentration via a sulfo-phospho-vanillin assay (SPVA) and interpolated from an E.coli calibrant (0, 50, 300, 500, and 1000 μg/mL) standard curve. After the lipid assay successfully passed, 5 μl SPLASH (equiSPLASH, Avanti Lipids) was added to each sample. These samples were then analyzed by LC-IMS-MS/MS on the timsTOF.

A pooled QC sample was prepared by combining 5 μl of each sample. Additional methanol with 10% chloroform was added for a total volume of 200 μl. This QC was then divided into 5 individual autosampler vessels so there would only be 1 injection/vial. The analytical samples were randomly analyzed by LC-IMS-MS/MS on the timsTOF, with the method blank being acquired first. A pooled QC was acquired before the first analytical sample, after the last analytical sample, and interspaced throughout the sequence to monitor instrument performance. Analytical sample loading was normalized for 1.5 μg total lipid content based on the SPVA lipid measurement. The LCMS method was programmed to allow a preceding 90/10 tune mix/sodium format plug prior to sample injection. This plug of calibrate allowed the data file to be re-calibrated (mass and IMS) prior to bioinformatics.

### Lipidomics Bioinformatics

Bruker, Compass, v.6.0 and Bruker, Metaboscape v.2022b were used for data filtration and lipid identification. The specific details can be found in the supplemental data. The data was first searched using rule-based IMS and/or MS/MS lipid matching. Annotations were then evaluated based on RT C# and H# trends. Remaining features were then searched using spectral (MSMS) matching from spectral lipid or metabolite libraries (hierarchical). MS1 only features were searched, but flagged with lower confidence.

### Lipidomics Response Analysis

Initially, the 88 peaks with positive LIPID MAPS^[14]^ annotations were analyzed using logistic GLM to identify species predictive of treatment group. The significant features were then used as anchors to validate analysis of partially annotated and un-annotated lipidomic features.

Analysis of the detected hits was designed around the truth table in Table S2 using the media control and the untreated bacteria conditioned media control to establish relevant contexts. For both presence and absence, the 0 or non-0 counts had to be consistent among all replicates. This led to a set of 6 groups with the most straight forward and confident being the set of molecules only present after treatment (hits that were not present in either control).

Statistical analysis was done prior to attempting identification of the molecules. The counts data for each experiment were normalized to the experimental count mean. Each group was analyzed by one-way ANOVA (α = 0.05) followed by Tukey’s Honest Significant Difference post hoc test to determine which groups differed significantly (p < 0.05) from one another. Hits with Tukey’s p < 0.05 were gathered in a subset for further dimensional reduction and identification. Dimensional reduction and clustering was accomplished using principal components analysis (PCA) and uniform manifold approximation and projection (UMAP^[15]^) via the python statsmodels and UMAP libraries.

### Candidate annotation of significant features

Beyond the 88 LIPID MAPS-annotated peaks, the full feature table comprised 15,757 molecular features across 32 injections. All differential and multivariate analyses were performed on the 26 biological samples; the 5 pooled-QC injections were used to assess analytical reproducibility (median feature %CV = 24.8%) and the single medium blank to seed the media/conditioned-media truth-table provenance classification. Per-contrast differences among treatment groups were evaluated by Welch’s t-test with Benjamini–Hochberg false-discovery-rate correction; features with q < 0.05 and |log_2_ fold-change| > 1 in any contrast were retained as the significant master set. Candidate identities were assigned to the significant features by accurate-mass matching against a theoretical bulk-lipid database spanning the major bacterial glycerophospholipid, glycerolipid, glycolipid, and fatty-acyl classes and five common adducts ([M+H]^+^, [M+NH_4_]^+^, [M+Na]^+^, [M+K]^+^, [M+H–H_2_O]^+^), retaining matches within ±0.005 Da and < 10 ppm and cross-checking molecular formulae against Bruker SmartFormula. Candidate identities remain at the accurate-mass / bulk-lipid-class level and were not confirmed by fragmentation. Matches were assigned to three confidence tiers: high-confidence annotations required agreement between the matched bulk-lipid formula and the independent SmartFormula elemental determination, with a subset additionally corroborated by a LIPID MAPS Structure Database entry of the same formula; medium-confidence annotations had a single unambiguous accurate-mass match within tolerance but lacked SmartFormula corroboration; and low-confidence annotations matched only at wider mass error (up to the 10 ppm ceiling) and are reported for completeness rather than as identifications. No tier distinguishes acyl-chain regiochemistry or resolves isobaric species sharing an ion formula (for example the FAHFA 34:1 / DG 31:0 pair), confidence refers to the reliability of the elemental composition and lipid class, not of a specific molecular structure.

## Results

### Collagen Microplate Assay

The initial collagen microplate biofilm assay demonstrated dose and compound-dependent reductions in established bacterial biofilms (Figure 1, Table S4). Pre-treatment Baseline cultures do not differ from 24hr incubated no-treatment cultures, indicating a mature, quiescent bacterial biofilm Notably, short lipid chain LEA-80 was the only effective compound in Gram-negative PAO-1, with no effect of longer chain LEA-160 or LEA-181, and vice-versa for Gram-positive *S. aureus*. This is consistent with prior screening in non-pathogenic Gram-negative *E. Coli* 29425, which is susceptible to LEA-80 treatment, but not LEA-160 or LEA-181 (data not shown).

**Figure 1.**
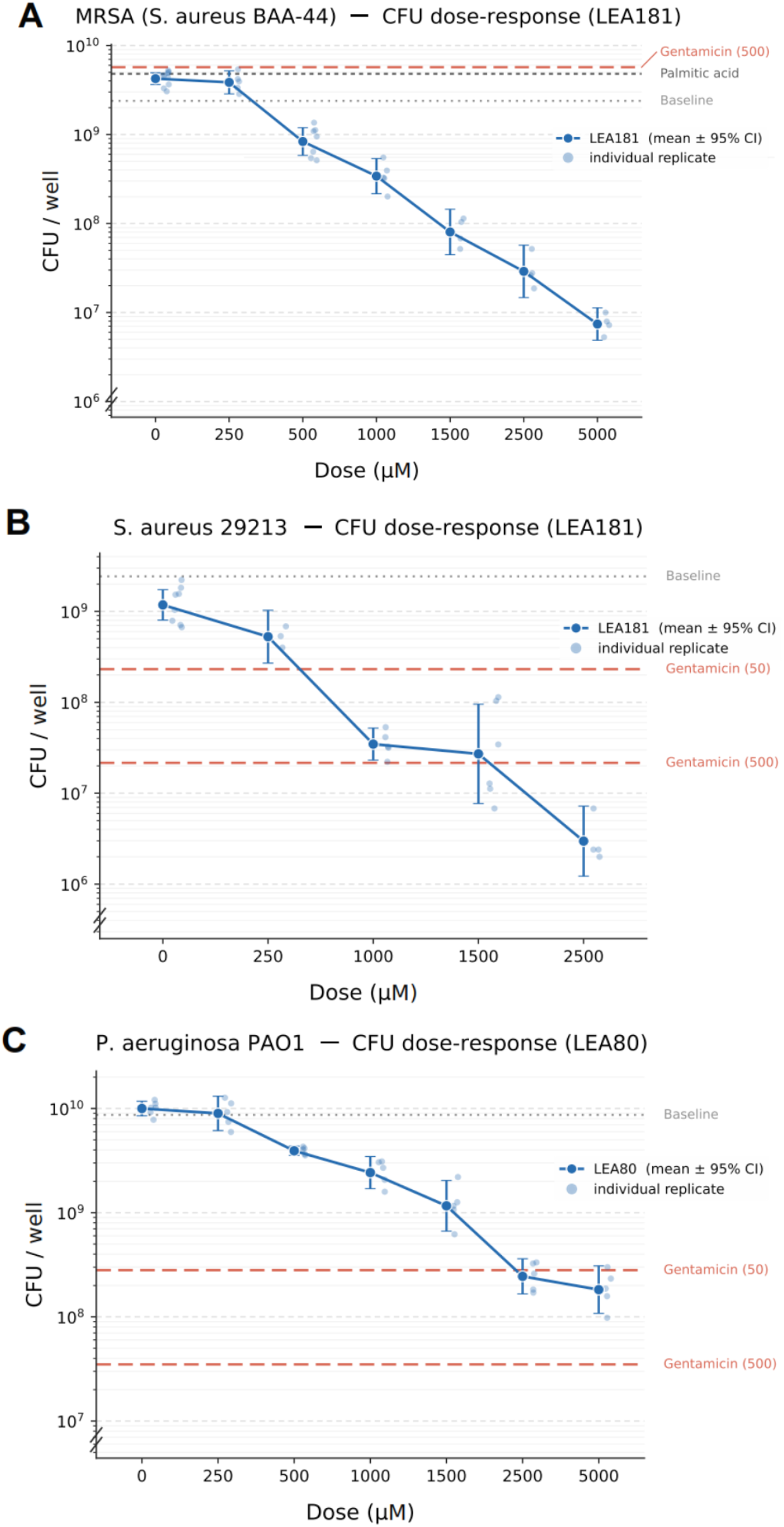
Collagen Microplate Assay Results. LEA treatment over 24hr produced dose-dependent CFU reductions in cultured biofilms of MRSA BAA-44 (A), *S. aureus* 29213 (B), and *P. aeruginosa* PAO1 (C). Mean pre-treatment biofilm CFU, Gentamicin treated (50μM, 500μM), and Palmitic acid treated (1mM) culture CFU are included as controls.

### Computational ligand-docking screen

In the global affinity model across all targets, LEAs, and endogenous ligands, GLM of binding affinities demonstrated significantly higher affinity of LEAs for the catalytic binding site of the phospholipase targets (-0.24, z=-5.14, 95% CI = [-0.47, -0.21]; Figure S1). In the per-ligand GLM model, arachidonic fatty acid analog LEA-204 was identified as the LEA analog with highest binding affinity (-0.65, z = -19.23, [-0.72, -0.59]) and synthesized.

### In-vitro Skin Construct Tolerability

Treatment of in vitro human skin constructs with topically formulated LEA compounds demonstrated no significant differences at any timepoint for LDH changes (Figure S2) and all timepoints were below the lower limit of detection for IL-1β (3.9 pg/mL). Histopathology of all skin constructs demonstrated epithelial regeneration of the biopsy punch simulated wound, but only LEA-181 treated explants were both completely closed by the 6-day timepoint (Figure S3).

### Minimal Inhibitory Concentration Studies

The initial work with the MIC assay in complete TSB media revealed that no LEA or organic fatty acid reached the MIC against *P. aeruginosa* (> 1.25mM), but LEA-181 and LEA-204 had some dose-dependent reduction in OD_600_ up through 1.25mM. All of the LEA’s reached MIC vs. MRSA and arachidonic acid also reached the MIC in the assay. The results are summarized in Table 1.

**Table 1.** Initial MIC results. LEAs consistently outperform their lipid-equivalent organic fatty acids. Only arachidonic acid had a measurable MIC for MRSA. Nothing impacted PAO1.

| Bacteria | Octanoic Acid | LEA-80 | Oleic Acid | LEA-181 | Arachidonic Acid | LEA-204 |
| --- | --- | --- | --- | --- | --- | --- |
| PAO1 | >180 μg/mL<br>(>1.25mM) | >352 μg/mL<br>(>1.25mM) | >352 μg/mL<br>(>1.25mM) | >520 μg/mL<br>(>1.25mM) | >380 μg/mL<br>(>1.25mM) | >552 μg/mL<br>(>1.25mM) |
| BAA44 | >180 μg/mL<br>(>1.25mM) | 88-176 μg/mL<br>(313-625μM) | >352 μg/mL<br>(>1.25mM) | <8 μg/mL<br>(<20μM) | 48 μg/mL<br>(156μM) | < 9 μg/mL<br>(<20μM) |

Due to the lack of activity in the MIC assay, no skins were inoculated with PAO1. After reaching a maximum CFU at Day 1 of growth, the biofilm CFU oscillated between 8-9 log_10_ CFU/mL. No treatment other than 10% bleach had a substantial impact on the viable biofilm bacteria (Figure S4).

Another MIC analysis was performed using a minimal essential media to test the MIC on BAA-44 and to generate materials for lipidomics analysis. The turbidimetric results can be found in Table 2 and minimal essential medium formulation is given in Supplementary Table S3.

**Table 2.**
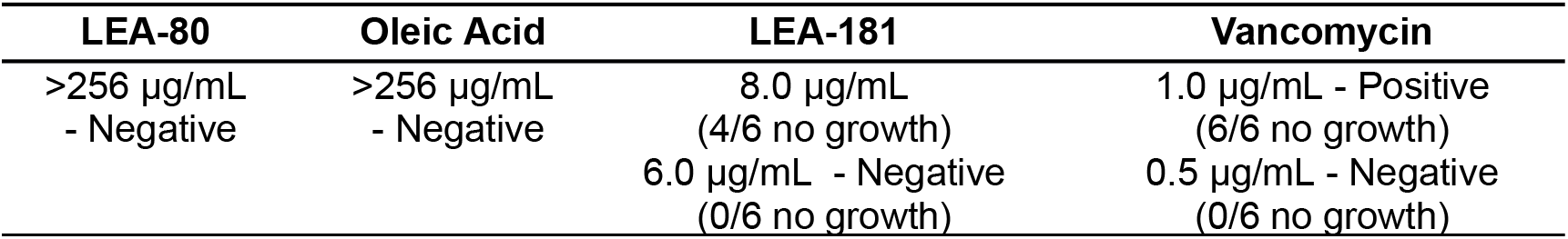
Results from the second MIC analysis.

| LEA-80 | Oleic Acid | LEA-181 | Vancomycin |
| --- | --- | --- | --- |
| >256 µg/mL<br>- Negative | >256 µg/mL<br>- Negative | 8.0 µg/mL<br>(4/6 no growth)<br>6.0 µg/mL - Negative<br>(0/6 no growth) | 1.0 µg/mL - Positive<br>(6/6 no growth)<br>0.5 µg/mL - Negative<br>(0/6 no growth) |

In minimal essential media, BAA-44 is sensitive to Vancomycin at 1.0 μg/mL but not 0.5 μg/mL. Of the test agents, only LEA-181 had an MIC within the range of the assay. Treatment of pig explant biofilms with non-formulated simple solutions did not reduce bacterial CFU, but in the ISO standardized assay effects were seen that were not attributable to the alkyl chain. Additional details were sought via a lipidomic study with bacteria in minimal media.

### Analysis of LIPID MAPS-annotated LC-MS/MS metabolomic data

The provenance and multivariate structure of the detected features are summarised in Supplementary Figures S5 and S7. GLM analysis of LIPID MAPS annotated lipidomic liquid chromatography/mass spectrometry sample data and treatment groups identified Phosphatidyl Glycerol 32:0 (PG 32:0) and Ricinoleic acid-C16:0 as statistically significant for treatment group assignment, at p<0.05. In *S. aureus* treated with LEA-181, there were dose dependent and statistically significant increases in peak intensity of PG 32:0 vs control, t = 3.02, 95% confidence interval for difference 728.02 to 17249.98 for 6μg LEA-181 treated samples and t = 3.21, 95% CI 626.41 to 4149.59 for 8μg/mL LEA-181 using unpaired t-test. Treatment with the natural fatty acid analog of LEA-181, oleic acid, demonstrated no difference in PG 32:0, instead significantly decreasing Ricinoleic acid-C16:0 vs control, t = 11.33, 95% confidence interval for difference -7462.56 to -4979.44. The widespread antibiotic vancomycin positive control demonstrated no changes in lipid composition. These are significant findings, validating a unique phospholipid anti-metabolite activity for LEA compounds and establishing a method for robust evaluation of bacterial lipid content for future drug development.

### Analysis of un-annotated LC-MS/MS features

An unsupervised overview of the full lipidomic feature table placed the treatment groups in a structured ordination. Principal-component analysis of the 522 ANOVA-significant features (26 samples) captured most of the variance in the first three components (PC1 43.5%, PC2 18.5%, PC3 12.7%; 74.7% cumulative). The untreated Control, oleic-acid, and low-dose LEA-181 (6 μg/mL) samples clustered together at the centre of the ordination, indicating that neither the fatty-acid vehicle nor the sub-inhibitory LEA-181 dose grossly reshaped the lipidome. The three perturbing conditions separated along distinct axes: LEA-80 was displaced strongly on PC2 (centroid +26.9 versus −4.2 for Control), the high-dose LEA-181 (8 μg/mL) samples were the extreme outliers on PC1 (centroid −45.7), and the vancomycin control shifted modestly away from Control on PC1. This separation establishes that the LEA compounds impose a reproducible, dose-scaled lipidomic perturbation distinct from both the untreated and fatty-acid-only states, and motivates the feature-level analysis that follows (Figure 2).

**Figure 2.**
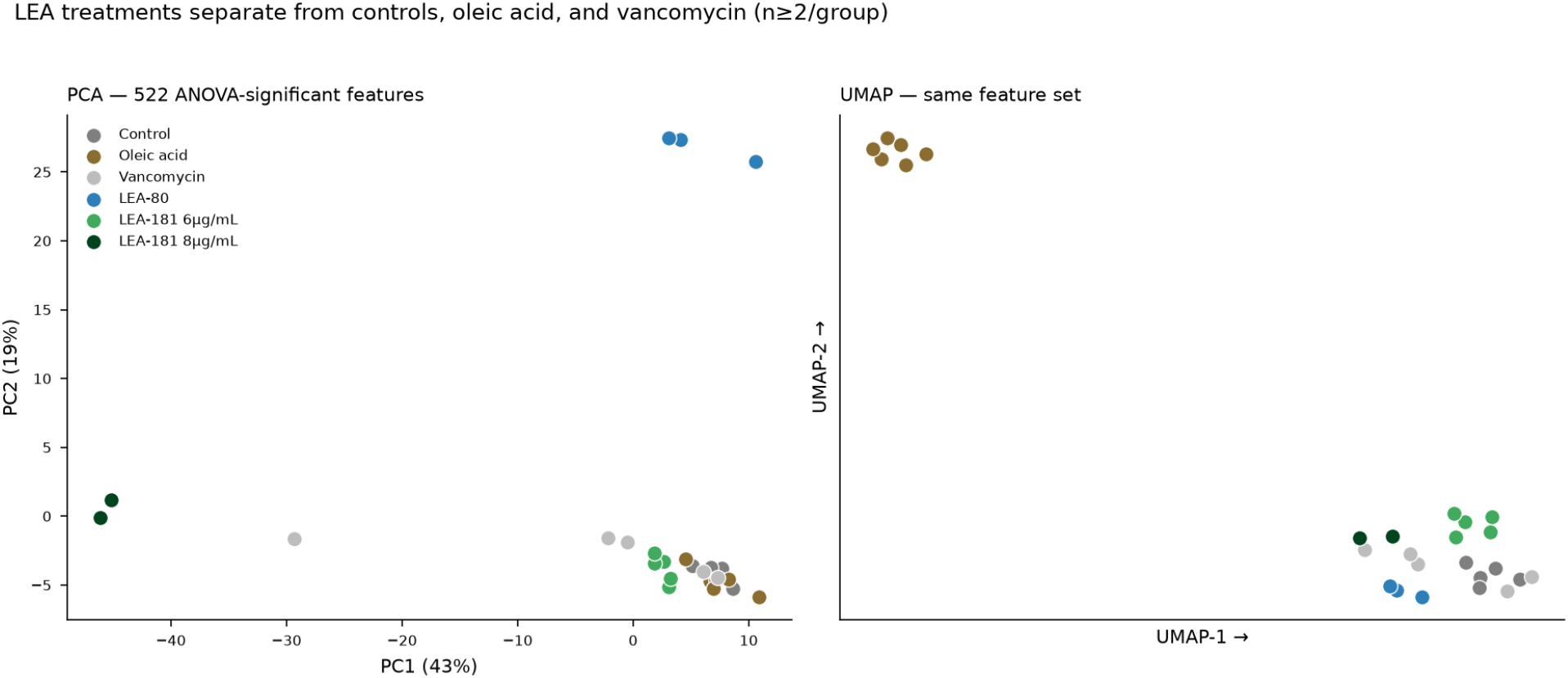
Ordination of the lipidomic profiles. Principal-component analysis (left) and UMAP (right) of the 522 ANOVA-significant features across 26 samples, colored by treatment group. Control, oleic acid, and low-dose LEA-181 overlap centrally, while LEA-80, high-dose LEA-181, and vancomycin separate along distinct axes.

The media/conditioned-media truth table (Table S2) partitioned the 15,757 features into media-derived (11,814), metabolized-media (840), bacteria-derived (691), and treatment-induced (164, of which 130 carried no prior annotation) sets. Across treatment groups, one-way ANOVA identified 522 features as significant for group assignment (408 previously un-annotated). The per-contrast significance filter yielded a master set of 1,918 pair-wise significant features, with the most significant hits in LEA-181 8μg/mL vs Control (866 hits) and LEA-181 8μg/mL vs oleic acid (801 hits). Of 1,304 de-duplicated significant features, 394 received a candidate class-level annotation (93 high-, 245 medium-, and 56 low-confidence), 261 of which were previously un-annotated. The two anchor species were recovered within tolerance: PG 32:0 as its ammonium adduct (−0.5 ppm) and the ricinoleic-acid FAHFA as its protonated ion (−0.3 ppm). The LEA-181 drug ion (behenoyl-ethanolamide, SPB 24:1;2O) was recovered among the treatment-induced features, accumulating strongly in both LEA-treated groups (e.g. +19 log_2_ versus control, q = 1.4×10^−6^). All identities are at the accurate-mass / bulk-lipid-class level and were not confirmed by fragmentation.

The distribution of annotations across lipid classes is shown in Figure S6. The three confidence tiers reflect the strength of orthogonal support for each accurate-mass assignment. The 93 high-confidence annotations combined a matched bulk-lipid formula with agreeing SmartFormula elemental determinations at a median mass error of 1.6 ppm, and 39 were additionally corroborated by a LIPID MAPS Structure Database entry of the same formula. The 245 medium-confidence annotations rested on a single unambiguous accurate-mass match (median 1.0 ppm) without confirmatory SmartFormula agreement, and the 56 low-confidence annotations matched only at wider mass error (median 5.6 ppm). Because no MS/MS was acquired, all tiers are class-level assignments and none resolves acyl-chain regiochemistry or isobaric species sharing an ion formula (for example the FAHFA 34:1 / DG 31:0 pair).

At the class level (confident annotations only), the two LEA compounds produced distinct signatures. Combining the LEA-181 doses (6 and 8 μg/mL, n = 7), phosphatidylglycerol was the single membrane-lipid class preserved or increased relative to control (median +0.21 log2; PG 32:0 +11.5 versus oleic acid, q = 0.037), whereas the surrounding classes fell: phosphatidic acid -0.51, lysyl-PG -0.56, diacylglycerol -0.62, triacylglycerol -0.99, and free fatty acids -0.56. This suppression deepened at the 8 μg/mL dose for every class except PG, which held near zero. LEA-80, by contrast, broadly lowered all membrane-lipid classes including PG (-0.32), and neither oleic acid nor the vancomycin control shifted the lipid profile. The class-level response and the corresponding pathway assignments are summarized in Figures 3 and 4.

**Figure 3.**
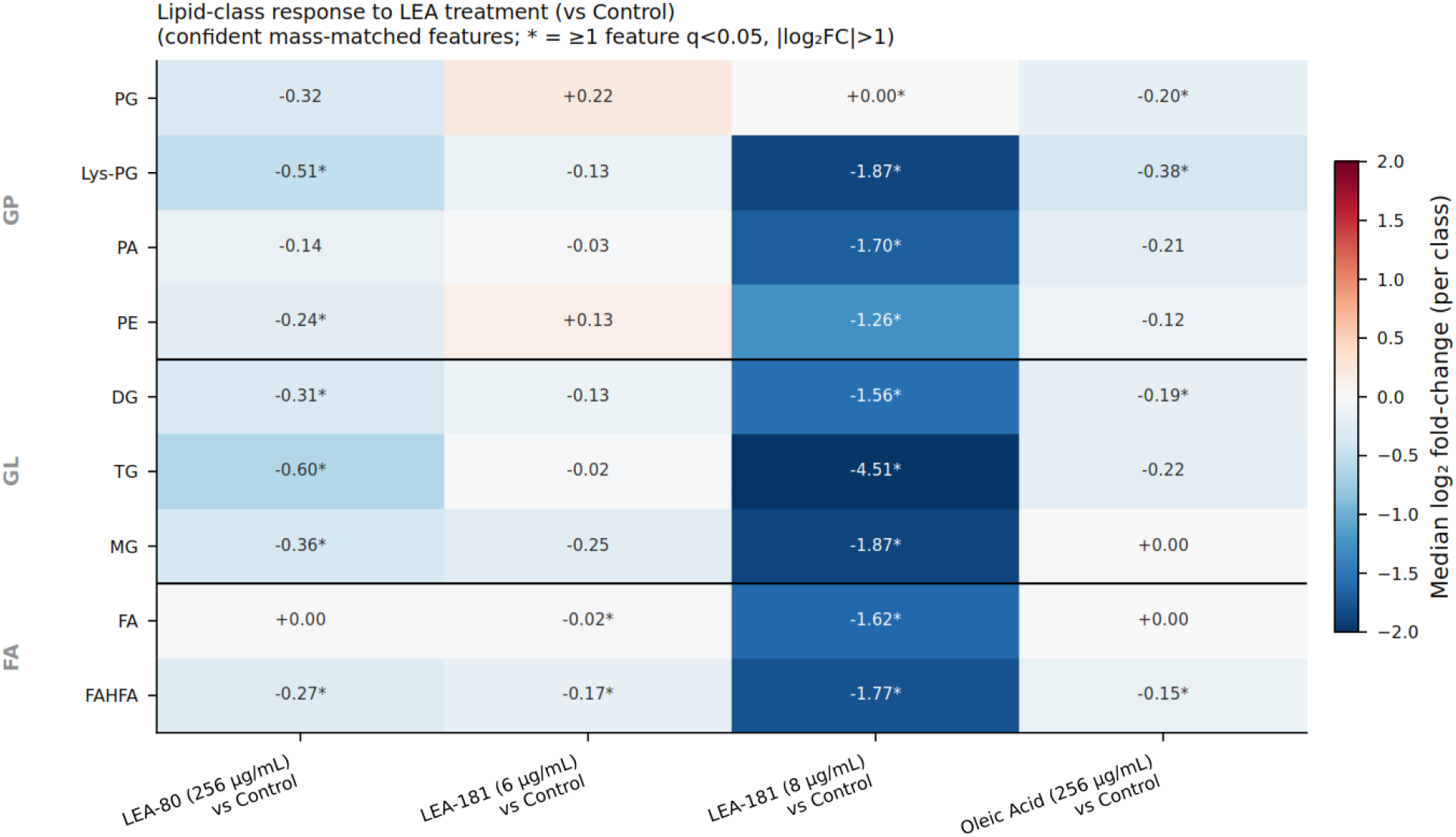
Lipid-class response to LEA treatment. Phosphatidylglycerol (PG) is preserved while surrounding classes fall. Median log_2_ fold-change per lipid class for each contrast; Control (n=5), LEA-80 (n=3) & Oleic Acid (n=6) at 256 μg/mL, LEA-181 6 μg/mL (n=5), 8 μg/mL (n=2). Lys-PG – Lysyl-phosphatidylglycerol, PA – Phosphatidic Acid, PE-Phosphatidylethanolamine, DG – Diacylglycerol, TG – Triacylglycerol, MG – Monoacylglycerol, GP – Glycerophospholipids, GL – Glycerolipids, FA – Fatty acid, FAHFA – Fatty acid ester of hydroxy fatty acid.

**Figure 4.**
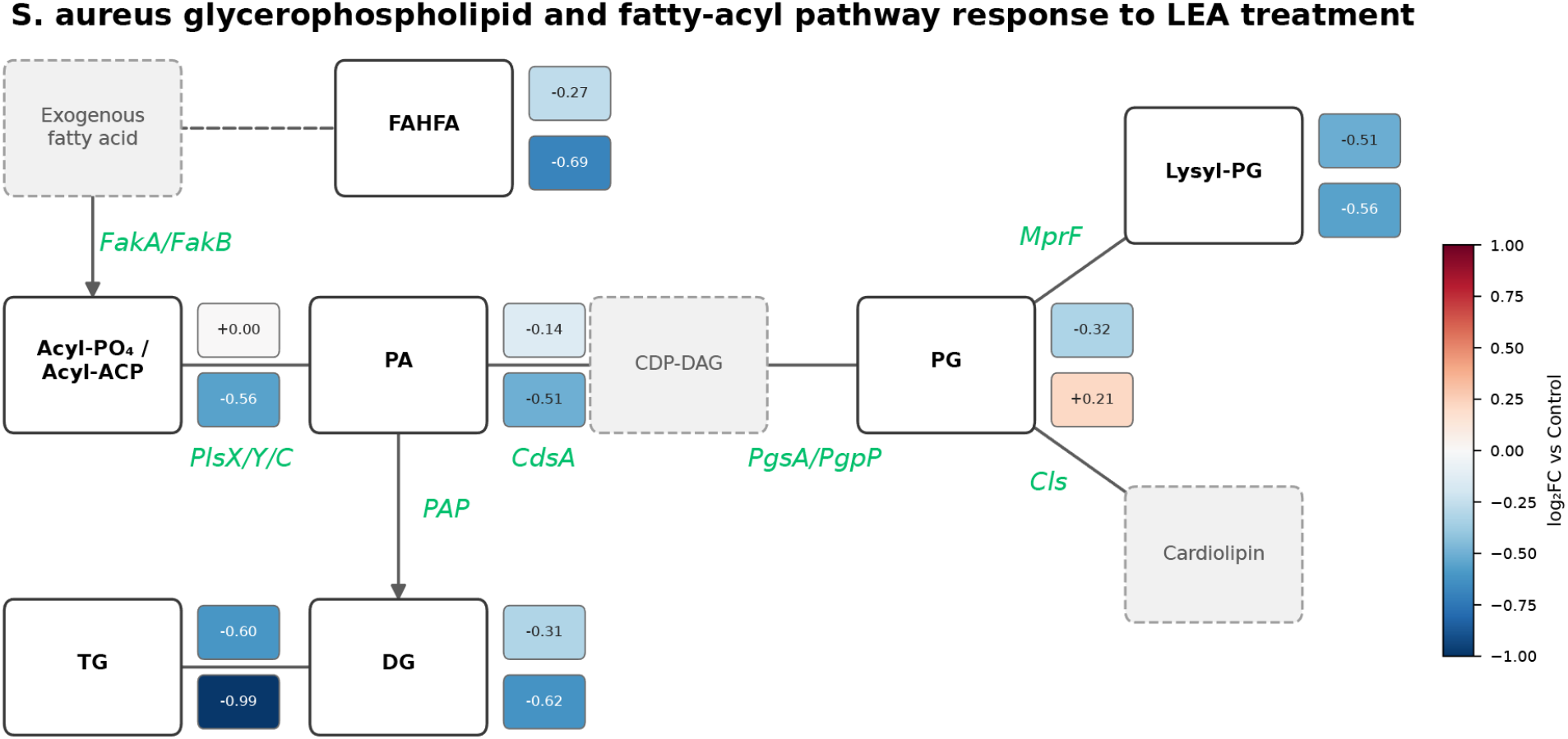
Lipid metabolic pathway of *S. aureus* with the observed treatment response. Nodes are colored by median class log_2_ fold-change, pooled 6 μg/mL and 8 μg/mL LEA-181, 256 μg/mL LEA-80 vs Control. Enzymatic steps (*PlsX/Y/C, CdsA, PgsA, MprF, Cls*) connect the glycerophospholipid classes and a fatty-acyl branch (FAHFA) is shown via a dashed edge to denote that the FAHFA-synthesizing route in *S. aureus* is not biochemically defined. The PG node is selectively maintained under LEA-181 relative to the surrounding suppressed classes.

## Discussion

### Summary of Principal Findings

This study establishes that lipid ether amines (LEAs) possess antibacterial activity against methicillin-resistant Staphylococcus aureus (MRSA) and that this activity depends on the cationic amine headgroup rather than on the alkyl chain alone. In a matched series in which each LEA was compared with its structurally equivalent fatty acid, the LEA consistently outperformed the corresponding fatty acid, demonstrating that the ether-linked amine contributes function beyond that of the lipid tail. Three convergent lines of evidence support an antibacterial role: reproducible minimum inhibitory concentrations (MICs) in minimal essential medium, dose-dependent reductions in MRSA biofilm CFU in the collagen microplate assay, and a treatment-specific lipidomic signature centred on the phosphatidylglycerol (PG) node. These gains were not reproduced against mature biofilms on porcine explants, a discrepancy we consider below. Taken together, the data position LEAs as a mechanistically distinct class of antimicrobials that act through bacterial membrane-lipid metabolism rather than through a single physicochemical membrane-lytic effect.

### Structure-Activity Relationships: The Amine Moiety Is Essential

The paired LEA/fatty-acid comparisons define the structural basis of activity. LEA-160 and LEA-181 reached an MIC and reduced MRSA biofilm CFU, whereas their matched fatty acids, palmitic and oleic acid, were inactive under the same conditions. The one exception among the fatty acids was arachidonic acid, which reached an MIC against MRSA. This is consistent with the established, chain-length- and unsaturation-dependent activity of free fatty acids, in which polyunsaturated species disrupt bacterial membranes directly while shorter saturated or monounsaturated chains do not.^[9, 16]^ That arachidonic acid was active while palmitic and oleic acid were not therefore reflects a property of the lipid tail itself and does not diminish the central observation: in every matched pair, the LEA was active where the fatty acid was not.

### Species-Specific Susceptibility Patterns

The compounds showed clear species selectivity. LEAs reached an MIC against MRSA but not against *Pseudomonas aeruginosa* PAO1 in the standard planktonic assay, in keeping with the additional permeability barrier imposed by the Gram-negative outer membrane. This selectivity was not absolute in the biofilm state: the medium-chain LEA-80 reduced PAO1 biofilm CFU in the collagen microplate assay despite failing to inhibit planktonic PAO1. The less metabolically active biofilm-associated bacteria^[1]^ may expose dependencies in lipid metabolism that are not accessible in rapidly dividing planktonic cells, offering a plausible route by which a compound inactive against planktonic Pseudomonas can still reduce its biofilm burden. Such state- and species-dependent behaviour is consistent with the distinct biofilm architectures that *P. aeruginosa* and *S. aureus* form on skin substrates.^[13]^

### Biofilm Efficacy: Microplate vs. Ex Vivo Models

In the collagen microplate, LEAs reduced biofilm CFU in a dose-dependent manner. On porcine explants, non-formulated aqueous LEA solutions, gentamicin, or vancomycin all failed to reduce mature biofilm. This pattern is expected from the biofilm literature: mature biofilms on biologically relevant substrates are markedly more tolerant to antimicrobials than biofilms on abiotic surfaces, with tolerance of roughly two orders of magnitude over planktonic cells and incomplete eradication even by chlorine-based disinfectants.^[1, 13]^ The negative explant result may also reflect a delivery limitation, since simple aqueous solutions provide none of the penetration or retention that an optimized formulation would. The favorable behavior of nanoparticle-formulated LEAs in the tolerability work supports pursuing engineered delivery.

### Lipidomics Evidence for Membrane Lipid Metabolism Disruption

The un-annotated feature mining reinforced and extended the annotated result. Sorting the full feature set with a media and conditioned-media truth table isolated a treatment-induced fraction and, after accurate-mass annotation, resolved a class-level picture that distinguishes the two active compounds. LEA-181 did not simply lower membrane lipids: it selectively preserved phosphatidylglycerol (most conspicuously PG 32:0), while phosphatidic acid, diacylglycerol, triacylglycerol, free fatty acids, and, notably, lysyl-PG all declined, an imbalance that widened with dose. LEA-80 instead suppressed all membrane-lipid classes, PG included. That PG selectivity is the feature that separates the more potent LEA-181 from LEA-80, and it converges on a single enzymatic step: the MprF-catalysed conversion of PG to lysyl-PG.

This signature is notable in light of recent *S. aureus* lipidomics, which found that the membrane species most distinguishing methicillin-resistant from methicillin-sensitive clinical strains are lysyl-PG and a diglycosyldiacylglycerol, placing the PG/lysyl-PG axis at the centre of the resistant phenotype.^[18]^ Mapping the *S. aureus* lipidome *in situ* during infection demonstrates that PG 32:0 is the most abundant bacterium-specific lipid within staphylococcal abscess communities and that ether and arachidonoyl lipids are dynamically remodelled, with MprF-mediated lysyl-PG modification a central resistance strategy.^[19]^ Our anchor species and the LEA-181 signature align with independently reported *in vivo* markers: LEA-181 accumulates the dominant in-situ bacterial lipid PG 32:0, while draining the lysyl-PG pool that shields the membrane from cationic attack.

Beyond the glycerophospholipid backbone, the fatty-acyl branch also responded to treatment. Fatty acid esters of hydroxy fatty acids (FAHFAs), including the confirmed ricinoleic-acid-C16:0 anchor species, were suppressed by both compounds, with median class log_2_ fold-changes of -0.27 under LEA-80 and -0.69 under the combined LEA-181 doses relative to Control. Unlike the PG node, which LEA-181 selectively preserved, the FAHFA branch fell alongside the free fatty-acid, diacylglycerol, and triacylglycerol pools, indicating a general drain on fatty-acyl supply rather than a PG-specific effect. Their coordinated decline is consistent with reduced fatty-acyl availability under both LEAs and distinguishes the broad lipid suppression common to the two compounds from the PG-selective effect that is unique to LEA-181.

### Relationship to Existing Lipid-Targeting Antibiotics

Among approved lipid-membrane-targeting antibiotics, daptomycin and the polymyxins are the closest comparators, and both are undermined by dedicated resistance mechanisms.^[10, 11]^ In *S. aureus*, the multiple peptide resistance factor MprF aminoacylates PG to lysyl-PG, raising the net surface charge of the membrane and repelling cationic antimicrobials,^[17, 20]^ a strategy that is central to the resistant phenotype observed in situ^[19]^. LEAs are nonetheless mechanistically distinct from daptomycin and the polymyxins: rather than partitioning into and disrupting the intact membrane, they were designed to engage lipid-metabolic enzymes upstream of the membrane, a distinction that may limit cross-resistance with membrane-lytic agents. The lipidomic signature makes a specific, testable prediction about how the LEAs engage this pathway. A cationic aminolipid that occupies the MprF active site or its lysyl-PG flippase channel would produce exactly the pattern observed: accumulation of the PG substrate together with depletion of the lysyl-PG product.^[17, 20-23]^ Reduced PG and elevated lysyl-PG accompanies daptomycin resistance in *S. aureus*,^[21]^ so the LEA-181 signature is that of a sensitizing rather than a resistance-promoting shift.

### Skin Tolerability and Potential for Topical Application

The tolerability data support a topical application. Treated human skin constructs showed no significant lactate dehydrogenase release and interleukin-1β below the limit of detection, indicating neither overt cytotoxicity nor a pro-inflammatory response at the concentrations tested. LEA-181-treated constructs additionally closed completely by the six-day timepoint, raising the possibility of a dual antimicrobial and wound-healing benefit. This combination is directly relevant to the clinical setting that motivates the work, in which biofilm-associated infections of chronic wounds and implants are the principal targets;^[3, 13]^ a topical agent that both suppresses biofilm and promotes closure would address an unmet need in that setting.

### Limitations

Several limitations should be borne in mind when interpreting these findings. The lipidomic analysis was acquired only in positive ion mode, which under-samples the anionic phospholipid classes and therefore leaves part of the membrane lipidome incomplete, including species central to the mechanism we propose. Compounding this, the candidate annotations of the un-annotated features rest on accurate mass and molecular formula at the bulk-lipid-class level and were not confirmed by MS/MS fragmentation. The specific acyl compositions, together with the PG and lysyl-PG assignments on which our mechanistic interpretation depends, will require validation against authentic standards. The antibacterial characterization is also limited. Only two bacterial species were tested, so broader-spectrum studies are needed to define the true activity range. Time-kill kinetics were not performed, leaving the bactericidal versus bacteriostatic nature of LEA activity unresolved. The collagen microplate model, although well suited to screening, does not fully reproduce the structural and chemical complexity of biofilms in vivo. Finally, sample sizes were modest throughout (n = 6–8 per group), and several dose-response estimates for biofilm reduction have wide confidence intervals (e.g. ED99 for LEA-160 against BAA-44 of 2,755 to 39,757μM).

### Future Directions

Several lines of work follow directly from these results. The most immediate is to test formulated LEA delivery, building on the nanoparticle approach used in the tolerability studies, against mature biofilm on the porcine explant model. This would establish whether the activity seen in the collagen-microplate assay translates, given more favorable exposure and pharmacokinetics vs an aqueous solution. In parallel, the antibacterial spectrum warrants extension to additional MRSA isolates and to other Gram-positive, biofilm-forming pathogens. Resolving the mechanistic questions raised here will require targeted lipidomics in negative-ion mode, both to capture the anionic phospholipid classes that positive-mode acquisition under-samples, lysyl-PG and cardiolipin among them, and to confirm the PG-to-lysyl-PG shift against authentic standards. Protein-ligand docking of the LEAs against MprF will provide a first test of the specific mechanism implied by the lipidomic data. Ultimately, efficacy against MprF-overexpressing and daptomycin-resistant strains, together with combinations against and together with conventional and cationic antimicrobials, will determine whether LEAs can act as sensitizers of the membrane-charge resistance node.

## Conclusions

Lipid ether amines constitute a structurally novel class of antimicrobials with activity against MRSA that derives from both the cationic amine headgroup and the lipid tail. Convergent MIC, biofilm, and lipidomic evidence indicates that their action is directed at membrane-lipid metabolism. The lipidomic signature of the most potent compound, LEA-181, selective preservation of phosphatidylglycerol with depletion of lysyl-PG and the surrounding lipid classes, implicates the MprF-catalysed PG-to-lysyl-PG step, which governs *S. aureus* resistance to cationic antimicrobials. Whether this reflects direct inhibition of MprF or a broader anti-metabolite effect on lipid supply remains to be established, and the biophysical and structural experiments outlined above will resolve it. By acting on lipid metabolism rather than the intact membrane, LEAs offer a differentiated and formulation-dependent strategy against biofilm-associated infection that merits further development.

## Supporting Information

The following supporting information is available: detailed LC-MS/MS acquisition, deconvolution, spectral-library search parameters, and data analysis; Table S1, VINA computational ligand docking screen targets; Figure S1, VINA computational ligand docking affinities, test compounds vs endogenous ligands; Figure S2, lactate dehydrogenase release from LEA-treated human skin constructs; Figure S3, histopathology of LEA-treated human skin constructs; Figure S4, Biofilm CFU post-treatment; Table S2, Lipidomic feature provenance truth-table; Table S3, Minimal essential media formulation; Table S4, Collagen microplate biofilm dose-response; Figure S5, Provenance classification of detected LC-MS/MS features, molecular weight (Da) versus retention time (min); Figure S6, Candidate lipid-class annotation coverage among significant features; Figure S7, UMAP ordination of the lipidomic feature space by provenance category.

## Author Contributions

D.J.G. and A.S.S. designed the study, performed experiments, analyzed data, and wrote the manuscript. All authors approved the final version.

## Notes

The authors declare the following competing financial interest(s): A.S. is affiliated with Integumed LLC and holds business and intellectual-property interests related to this work. D.J.G. declares no competing interest.

## Acknowledgements

Dr. Laura S. Bailey and the University of Florida Mass Spectrometry Research and Education Center for their support with obtaining and sharing the lipidomics data.

## Data Availability Statement

The whole lipidomic LC/MS-MS dataset will be published to The Metabolomics Workbench (https://www.metabolomicsworkbench.org/) and additional data generated and/or analyzed during the current study are available from the corresponding author upon reasonable request.

