## Supporting Information for "The Testing and Characterization of the Bacterial Response to Novel Lipid Ether Amine-Based Antibiotics"

*Liquid Chromatography*

Injection system: Bruker Elite series degasser, binary pump, column compartment, and autosampler
Injection volume: 2 µg lipid content; volume variable (3-6 µL, uL-pickup method, using 60/40 H2O/ACN for the pickup solvent)
QCs (uL-pickup method) – 5 µL, pooled sample
Blanks (uL-pickup method) – 10 µL, MeOH
Column: Waters Acquity UPLC CSH C18 (2.1 mm x 100 mm, 1.7 µm)
Column temperature: 55°C
Autosampler temperature: 4°C

*Mobile Phases*:
A = 60/40 acetonitrile/water + 10 mM ammonium formate + 0.1% formic acid
B = 90/8/2 isopropanol/acetonitrile/water + 10 mM ammonium formate + 0.1% formic acid

*Gradient(s) for LC-MS analysis:*

| Time (min) | 0 | 2 | 15 | 20 | 21 | 26 |
|---|---|---|---|---|---|---|
| %B | 50 | 50 | 98 | 98 | 50 | 50 |
| %A | 50 | 50 | 2 | 2 | 50 | 50 |

Flow rate = 0.4 mL/min

*Mass Spectrometry:*
Instrument: Bruker Daltonics, tims-tof pro II TIMS-QqTOF MS
Ionization: electrospray ionization (ESI) (analyzed in positive mode)
Wide range, mid-mass lipids optimized method
(*m/z* 100 – 2000; $k0^{-1}$ 0.55-1.92 V*s/$cm^2$)
Capillary voltage: 4.5 kV
Gas temperature: 220˚C
Drying gas ($N_2$): 10.0 L/min
Nebulizer: 2.2 bar
TIMS ramp time: 100.0 ms
MS2 parameters:
Activation: Collision induced dissociation (CID)
Parallel accumulation and serial fragmentation (PASEF)
PASEF MS/MS scans: 2
Inclusion IMS range: 0.55 – 1.90 V*s/$cm^2$
Charge state: 1
Active exclusion parameters:
if selected 1 times, released after 0.1 min,
reconsidered if In/I(n-1)>3.
Collision energy (CE)
Isolation: 100 – 1350 m/z +/- 2
1/k0: 0.55 – 1.90 V*s/cm2
Energy: 30 – 30 eV
CCS: 1.0 / 3.0 %

*Initial data filtering:*

| | |
|---|---|
| Signal threshold: | 500 counts |
| Minimum peak length: | 80 spectra |
| Recursive: | 50 spectra |
| Feature filtering: | 3/70 samples |
| Recursive: | 4/70 samples |
| Group filtering: | 60% grouped samples |
| Retention time range: | 0.9 – 14.5 min |
| Alignment window: | 25 s |
| Mass range: | m/z 200 – 2000 |
| Ion deconvolution | |
| Primary: | $+H]^+$ |
| Seed: | $+NH_4]^+$, $+Na]^+$, $+K]^+$ |
| Common: | $+H\text{-}H_2O]^+$ |

*Lipid rule-based IMS/MSMS searching parameters (narrow / wide):*

*Bruker lipid species annotation*

Search all categories and classes
Validation by MSMS required
Primary ion reassignment

| | |
|---|---|
| m/z matching: | 2.0 ppm / 5.0 ppm |
| mSigma matching: | 20 / 1000 |
| MS/MS scoring: | 900 / 500 |
| CCS: | 1.0 / 3.0 % |

*Spectral Library searching parameters (narrow / wide):*

LipidBlast v.2022
PNNL lipids positive
MSDIAL-Tandem Mass Spectral Atlas, vs66
Bruker MetaboBASE personal library v3.0
MassBank
LC-MS-MS Positive Mode
HCE cell lysate lipids
ECG acyl amides C4-C24
IMS oxidized phospholipids
HMDB metabolomics database
Birmingham UHPLC MS positive
GNPS natural products/metabolites (without propagation)
LipidFinder
LIPID MAPS

| | |
|---|---|
| m/z matching: | 2.0 ppm / 5.0 ppm |
| mSigma matching: | 20 / 1000 |
| MS/MS scoring: | 900 / 500 |

Data was first searched using rule-based IMS and/or MS/MS lipid matching. Annotations were then evaluated based on RT C# and H# trends. Remaining features were then searched using spectral (MSMS) matching from the above spectral lipid or metabolite libraries (hierarchal). MS1 only features were not searched.

### Supplementary Figures

| pdb_id | name | type | species | Endogenous Ligands | Search box |
|---|---|---|---|---|---|
| 5G3M | sPLA2 | Secreted Phospholipase A2 | *H. sapiens* | pc, pe, sm, ps | -2,-1,-1,20,30,30 |
| 2ZKM | PLC | Phospholipase C | *H. sapiens* | pi | 38,21,95,30,30,30 |
| 5JG8 | aSmase | Acid Sphingomyelinase | *H. sapiens* | sm | 5,40,43,14,14,14 |
| 5TCD | alk-Smase | Alkaline Sphingomyelinase | *H. sapiens* | sm | 38,49,54,30,30,20 |
| 5uvg | nSMase2 | Neutral Sphingomyelinase | *H. sapiens* | sm | 10,94,62,40,40,30 |
| 6OHO | PLD2 | Phospholipase D2 catalytic domain | *H. sapiens* | pc | 14,21,3,14,14,14 |
| 6OHR | PLD1 | Phospholipase D1 catalytic domain | *H. sapiens* | pc | 8,1,56,14,14,14 |
| 1KP4 | PLA2 | Surface-associated PLA2 | *S.violaceoruber* | pc, pe, sm, ps | 1,17,29,32,32,32 |
| 1AH7 | Bc-PLC | Phosphatidylcholine hydrolysing Phospholipase C | *B.cereus* | pc | 40,26,5,20,20,20 |
| 1ZWX | SmcL | Hemolytic sphingomyelinase C | *L.ivanovii* | sm | 17,25,84,32,32,28 |
| 2DDT | Bc-SMase | Sphingomyelinase C | *B.cereus* | sm | -4,-13,1,16,16,16 |
| 3I5V | Beta-toxin | Sphingomyelinase C | *S.aureus* | sm | 0,-3,-16,28,28,40 |
| 3tu3 | ExoU | Type III effector cytotoxin | *P.aeruginosa* | sm | 64,38,18,20,30,30 |
| 4as2 | PchP | Phosphorylcholine Phosphatase | *P.aeruginosa* | pc | 27,51,46,40,40,30 |
| 4F2T | Pi-PLC (SA) | 1-phosphatidylinositol phosphodiesterase | *S. aureus* | pi | -23,-3,3,20,30,30 |
| 4G33 | Lox | Phospholipid-Lipoxyg enase | *P. aeruginosa* | pc, pe, sm, ps | -27,-14,-21,30,30,40 |
| 5FYP | Pi-PLC (PA) | 1-phosphatidylinositol phosphodiesterase | *P. aeruginosa* | pi | 37,-7,-18,30,30,30 |

**Table S1.** Docking targets with refined catalytic binding sites. Crystallographic (PDB) structures of secreted bacterial virulence factors and endogenous human phospholipases used as receptors for AutoDock Vina docking of the LEA analogs, endogenous lipid ligands, and the propranolol decoy, with each target's endogenous ligands and search-box definition. These refined targets are a subset of the 34 structures screened in the full pipeline.

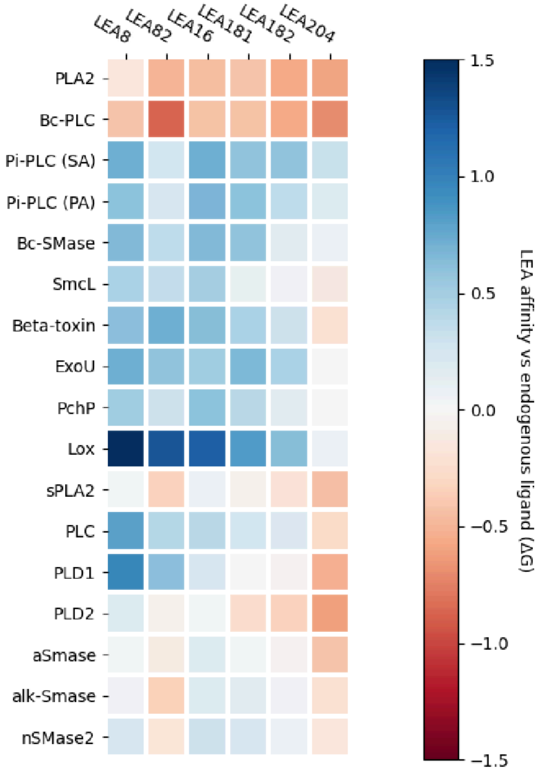


**Figure S1.** VINA docking study results. Heatmap values are mean deltaG (kcal/mol) for 8 replicates, endogenous ligands – compound, for targets in Table S1.

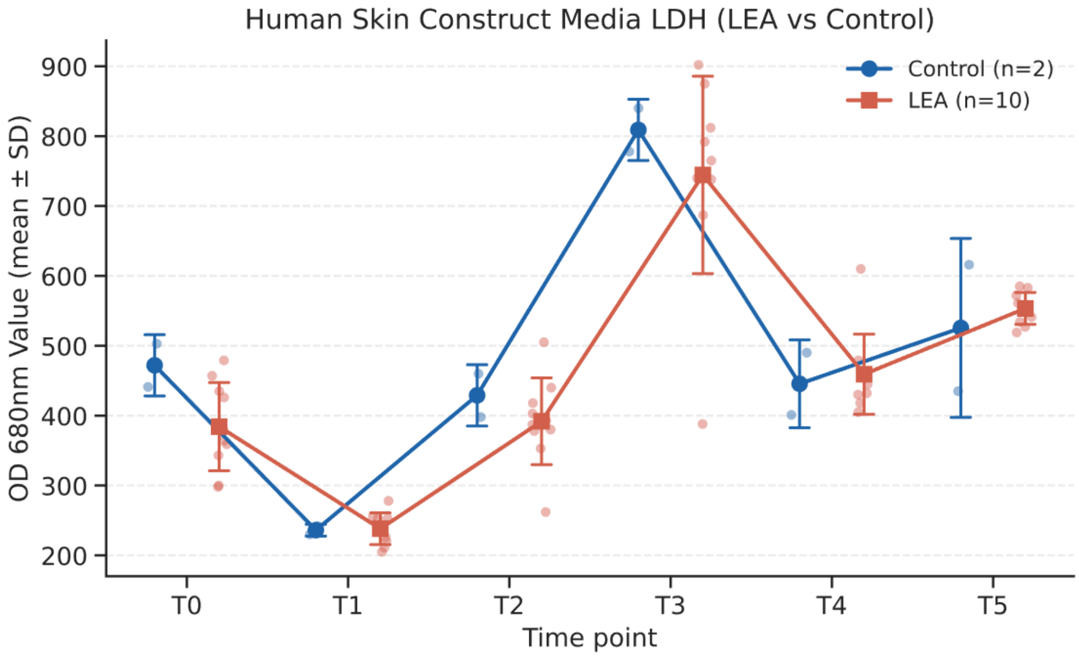


**Figure S2.** Lactate dehydrogenase release from LEA-treated human skin constructs. Media LDH (OD 680 nm, mean ± SD) over the treatment time course (T0–T5) for full-thickness in vitro human skin constructs (MatTek EpiDermFT) topically treated with LEA compounds (n = 10) versus vehicle control (n = 2). No significant difference in cytotoxicity was observed at any timepoint.

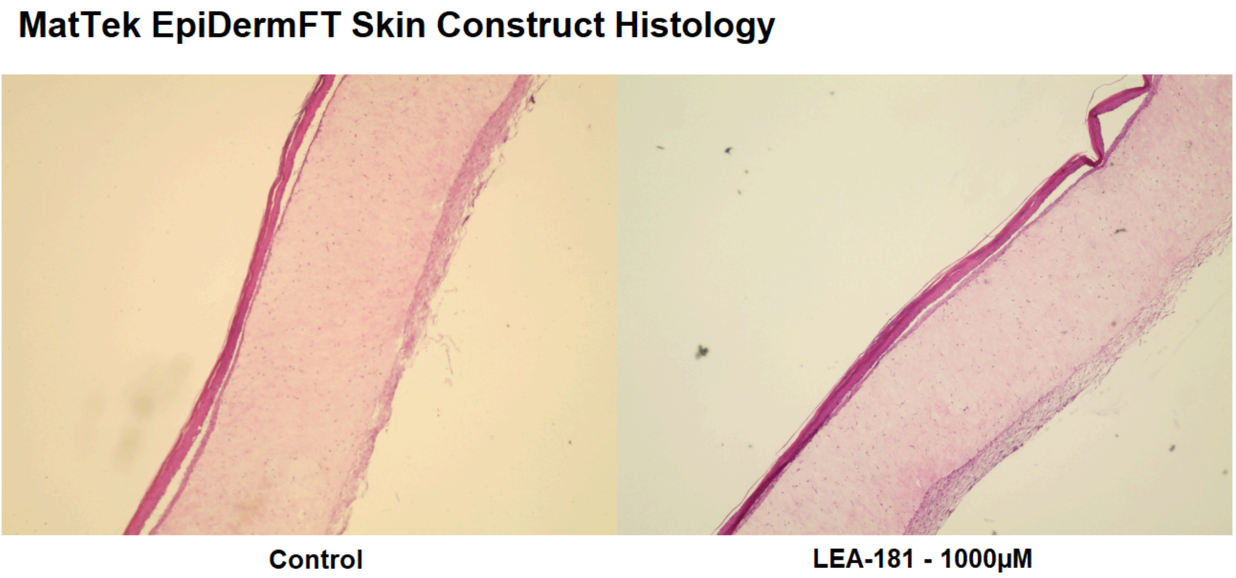


**Figure S3.** Histopathology of LEA-treated human skin constructs. Representative haematoxylin and eosin sections of biopsy-punch–wounded skin constructs after 6 days, comparing vehicle control (left) with LEA-181 at 1 mM (right). Both show epithelial regeneration and complete closure across the simulated wound.

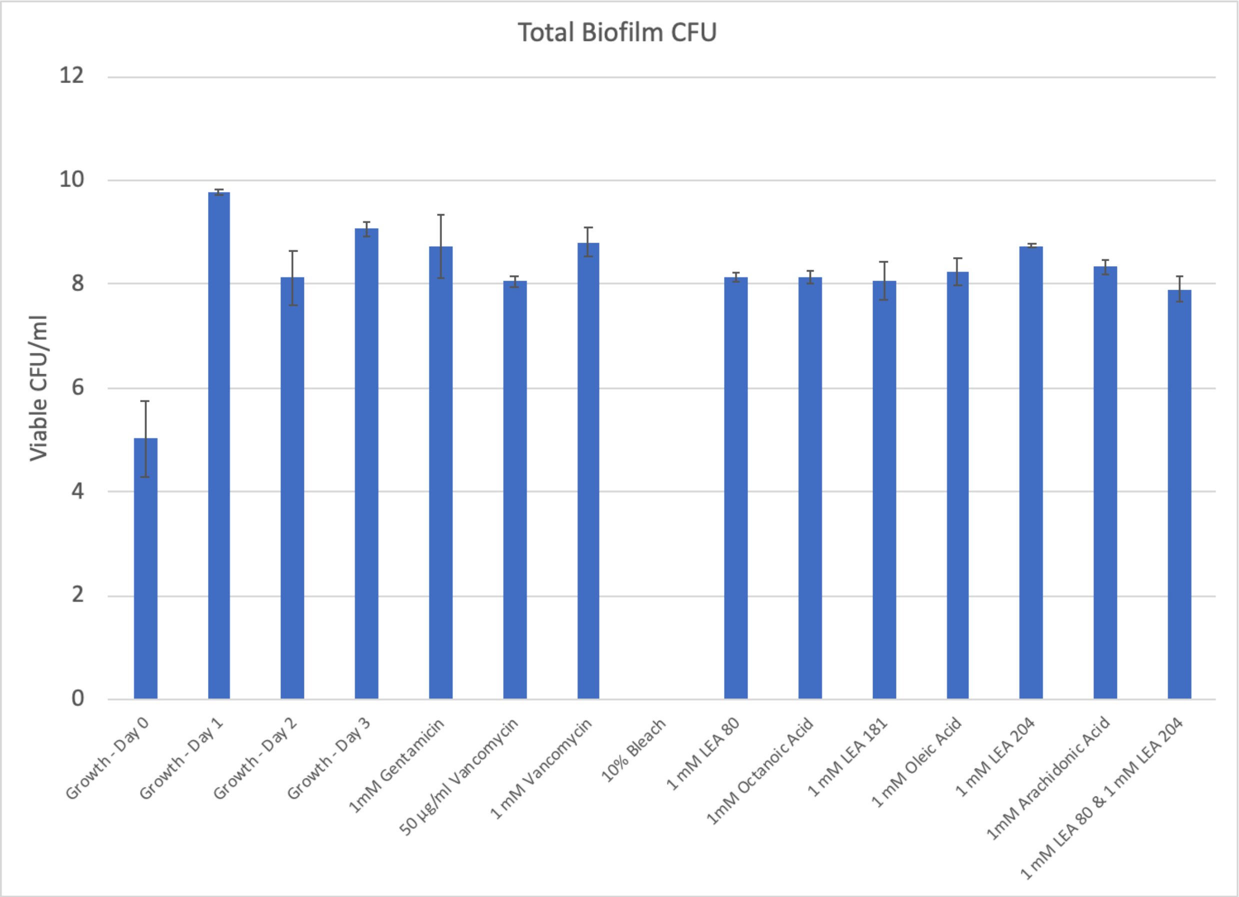


**Figure S4**. Biofilm CFU post-treatment. Only the bleach-treated samples had substantial reductions.

<table>
<tr><td colspan="2" rowspan="2"></td><td colspan="2">Fresh Media</td></tr>
<tr><td>Present</td><td>Absent</td></tr>
<tr><td rowspan="3">Conditioned Media</td><td>Present</td><td>Media-derived molecules<br><br>Significant Change ($p < 0.05$):<br>Metabolism altered<br>No Change ($p \geq 0.05$):<br>No effect</td><td>Media-derived molecules<br><br>Significant Change ($p < 0.05$):<br>Metabolism altered<br>No Change ($p \geq 0.05$):<br>No effect</td></tr>
<tr><td>Absent</td><td>Media-derived molecules which were metabolized away in the control<br><br>Present ($p < 0.05$):<br>Metabolism blocked<br>Absent:<br>No effect</td><td>Newly Synthesized molecules<br>$p < 0.05$:<br>1. New response to intervention<br>2. The test articles themselves<br>$p \geq 0.05$:<br>Noise</td></tr>
<tr><td>Inconsistent</td><td>Media-derived noise</td><td>Bacteria-derived noise</td></tr>
</table>

**Table S2.** Truth-table classification of detected LC-MS/MS features. Each feature was assigned to a provenance category from its presence or absence in fresh medium versus conditioned medium across all replicates, distinguishing media-derived, bacteria-derived, newly synthesized (treatment-induced), and inconsistent (noise) sets.

| **BAA-44 Minimal Essential Media** | |
|---|---|
| $KH_2PO_4$ | 3 g/L |
| $Na_2HPO_4$ | 12.8 g/L |
| NaCl | 0.5 g/L |
| $NH_4Cl$ | 0.5 g/L |
| $MgSO_4$ | 2mM |
| $CaCl_2$ | 0.1mM |
| Glucose | 1% |
| Casaminoacids | 1% |
| Thiamine-HCl | 1 mM |
| Nicotinamide | 0.05 mM |

**Table S3.** Minimal Essential Media formulation

| PAO1 | Dose | Mean | SD |
|---|---|---|---|
| Pre-Treatment | - | 9.94 | 0.04 |
| Gentamicin | 50 μM | 8.43 | 0.12 |
| Gentamicin | 500 μM | 7.54 | 0.37 |

| PAO1 + LEA-80 | Estimate (μM) | Lower | Upper |
|---|---|---|---|
| $ED_{50}$ | 443 | 362 | 543 |
| $ED_{90}$ | 990 | 590 | 1659 |
| $ED_{99}$ | 2377 | 911 | 6205 |

| MRSA BAA-44 | Dose | Mean | SD |
|---|---|---|---|
| Pre-Treatment | - | 9.31 | 0.20 |
| Palmitic Acid | 1000 μM | 9.68 | 0.04 |
| Gentamicin | 500 μM | 9.76 | 0.08 |

| BAA-44 + LEA-160 | Estimate (μM) | Lower | Upper |
|---|---|---|---|
| $ED_{50}$ | 306 | 203 | 460 |
| $ED_{90}$ | 1656 | 912 | 3007 |
| $ED_{99}$ | 10465 | 2755 | 39757 |

| BAA-44 + LEA-181 | Estimate (μM) | Lower | Upper |
|---|---|---|---|
| $ED_{50}$ | 338 | 196 | 583 |
| $ED_{90}$ | 684 | 178 | 2628 |
| $ED_{99}$ | 1873 | 153 | 22859 |

| SA 29213 | Dose | Mean | SD |
|---|---|---|---|
| Pre-Treatment | - | 9.36 | 0.21 |
| Gentamicin | 50 μM | 8.36 | 0.10 |
| Gentamicin | 500 μM | 7.32 | 0.11 |

| SA 29213 + LEA-160 | Estimate (μM) | Lower | Upper |
|---|---|---|---|
| $ED_{50}$ | 276 | 105 | 729 |
| $ED_{90}$ | 1532 | 434 | 5415 |
| $ED_{99}$ | 9951 | 477 | 207665 |

| SA 29213 + LEA-181 | Estimate (μM) | Lower | Upper |
|---|---|---|---|
| $ED_{50}$ | 212 | 101 | 447 |
| $ED_{90}$ | 622 | 121 | 3199 |
| $ED_{99}$ | 2009 | 49 | 82700 |

**Table S4.** Collagen microplate biofilm dose-response. Log-logistic effective-dose estimates (ED50/ED90/ED99, μM) and 95% confidence intervals for LEA compounds against *P. aeruginosa* PAO1, MRSA BAA-44, and *S. aureus* ATCC 29213, with pre-treatment and reference-agent (gentamicin, palmitic acid) controls; dose-response curves were fitted with the R drc package.

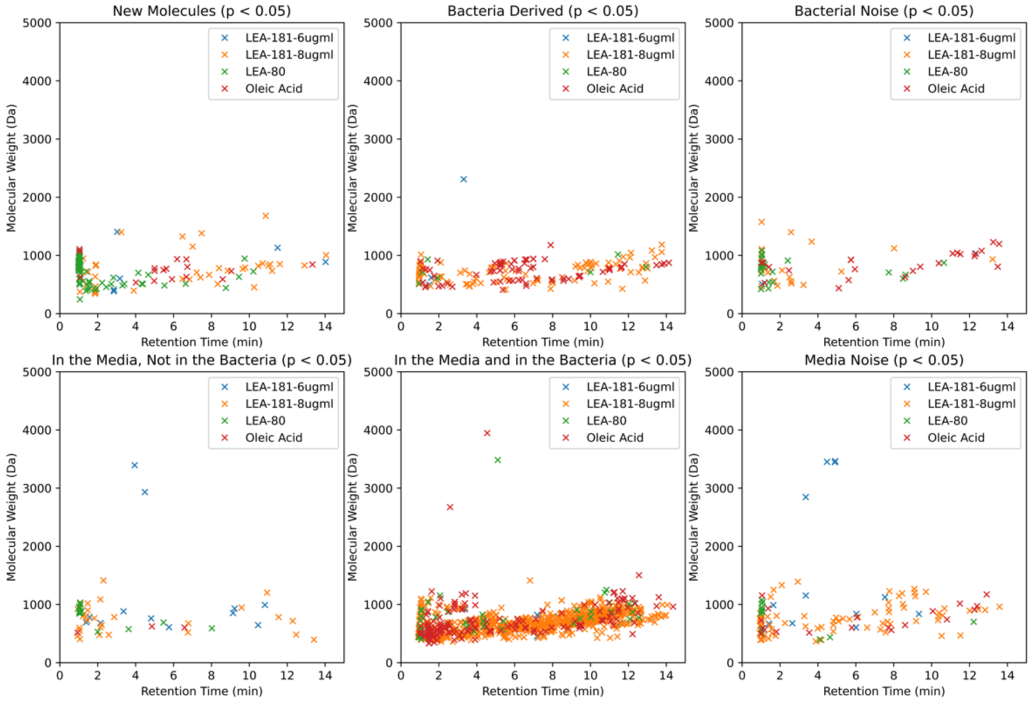


**Figure S5.** Provenance classification of detected LC-MS/MS features. Molecular weight (Da) versus retention time (min) for features passing significance ($p < 0.05$), partitioned by the media/conditioned-media truth table into six provenance categories — new molecules, bacteria-derived, bacterial noise, present in media but not bacteria, present in both media and bacteria, and media noise. Points are colored by treatment group (LEA-181 at 6 and 8 µg/mL, LEA-80, and oleic acid).

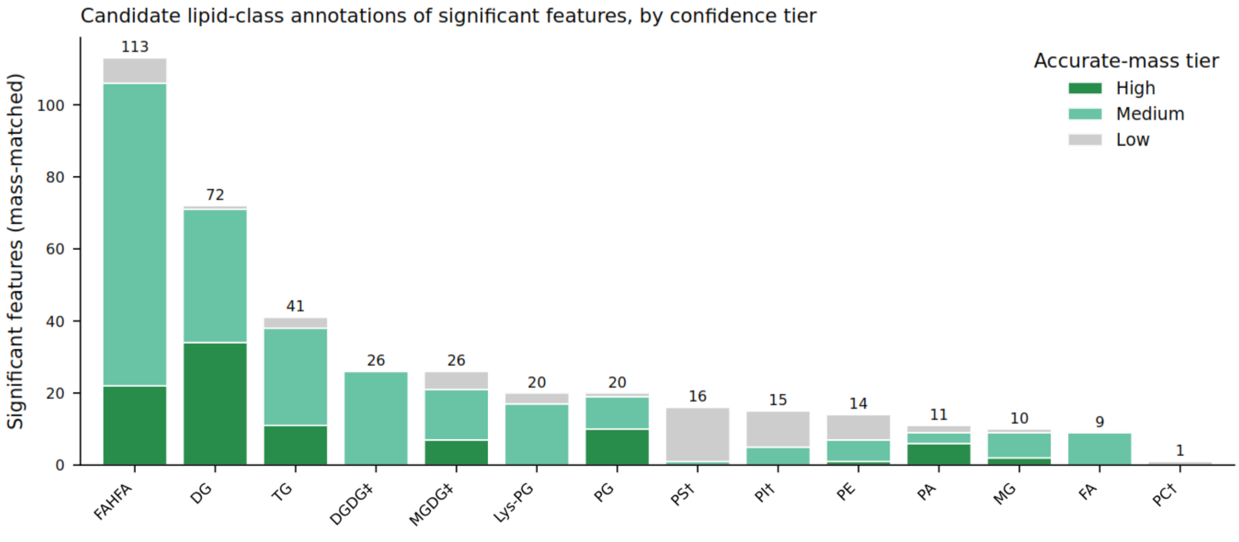


**Figure S6.** Candidate lipid-class annotation coverage among significant features. Counts of significant features receiving a candidate identity in each bulk-lipid class by accurate-mass matching (±0.005 Da, <10 ppm), split by confidence tier (high, medium, low). Identities are class-level accurate-mass assignments and were not confirmed by MS/MS.PS, PI, PC, DGDG, and MGDG are primarily media-derived and spurious matches, included for completeness.

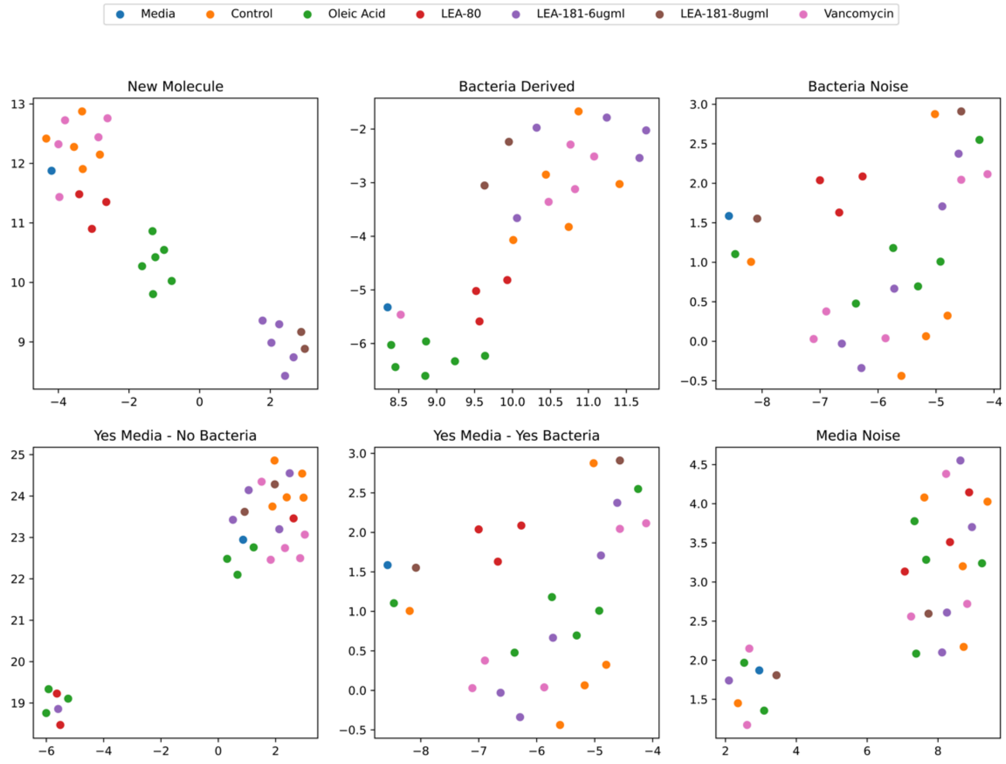


**Figure S7.** UMAP ordination of the lipidomic feature space by provenance category. Uniform manifold approximation and projection of the detected features, faceted by the same six truth-table provenance categories as in Figure S5 (new molecule, bacteria-derived, bacterial noise, media-not-bacteria, media-and-bacteria, media noise) and colored by treatment group (media, control, oleic acid, LEA-80, LEA-181 at 6 and 8 μg/mL, and vancomycin).
